# Discovering biomarkers associated and predicting cardiovascular disease with high accuracy using a novel nexus of machine learning techniques for precision medicine

**DOI:** 10.1101/2023.09.08.553995

**Authors:** William DeGroat, Habiba Abdelhalim, Kush Patel, Dinesh Mendhe, Saman Zeeshan, Zeeshan Ahmed

**Author notes:** **Corresponding author:** Zeeshan Ahmed, Rutgers Institute for Health, Health Care Policy and Aging Research, Rutgers University, 112 Paterson Street, New Brunswick, 08901, NJ, USA.

## Abstract

Personalized interventions are deemed vital given the intricate characteristics, advancement, inherent genetic composition, and diversity of cardiovascular diseases (CVDs). The appropriate utilization of artificial intelligence (AI) and machine learning (ML) methodologies can yield novel understandings of CVDs, enabling improved personalized treatments through predictive analysis and deep phenotyping. In this study, we proposed and employed a novel approach combining traditional statistics and a nexus of cutting-edge AI/ML techniques to identify significant biomarkers for our predictive engine by analyzing the complete transcriptome of CVD patients. After robust gene expression data pre-processing, we utilized three statistical tests (Pearson correlation, Chi-square test, and ANOVA) to assess the differences in transcriptomic expression and clinical characteristics between healthy individuals and CVD patients. Next, the Recursive Feature Elimination (RFE) classifier assigned rankings to transcriptomic features based on their relation to the case-control variable. The top ten percent of commonly observed significant biomarkers were evaluated using four unique ML classifiers (Random Forest, Support Vector Machine, Xtreme Gradient Boosting Decision Trees, and k-Nearest Neighbors). After optimizing hyperparameters, the ensembled models, which were implemented using a soft voting classifier, accurately differentiated between patients and healthy individuals. We have uncovered 18 transcriptomic biomarkers that are highly significant in the CVD population that were used to predict disease with up to 96% accuracy. Additionally, we cross-validated our results with clinical records collected from patients in our cohort. The identified biomarkers served as potential indicators for early detection of CVDs. With its successful implementation, our newly developed predictive engine provides a valuable framework for identifying patients with CVDs based on their biomarker profiles.

## Introduction

Artificial intelligence (AI) and machine learning (ML) encompasses a plethora of supervised and unsupervised methodologies for scrutinizing genomics data, culminating in the formation of multivariate statistical instruments [1]. The proficient implementation of AI/ML techniques holds the promise of fostering an augmented comprehension of diseases at the systemic level, unveiling the intricacies of genomic regulatory networks. By leveraging AI/ML approaches, clinical and genomics data can undergo statistical analysis and classification, enabling the prediction of high-risk patients. AI/ML can be deployed to capture genetic sequences associated with chronic diseases, categorize phenotypes based on knowledge about human diseases and establish population dimensions for rare diseases [1, 2]. Genetic studies have facilitated disease prognosis [3, 4], the identification of genetic regions and variants that influence disorders, and the functional assessment of these regions [5, 6, 7]. While holding great prospects, the formidable task at hand lies in analyzing the immense magnitude of recognized (and unrecognized) genetic variations and leveraging this knowledge to facilitate diagnosis, ascertain risk, and forecast treatment responses among heterogenous human populations [8]. This challenge is being addressed through precision medicine which encompasses the integration of clinical and genomics data to enable predictive treatment within a diverse cardiovascular disease (CVD) population [9]. The primary objective of personalized medicine is to analyze a patient’s genetic makeup to identify crucial biomarkers and enhance comprehension of the underlying pathophysiology of intricate disorders such as CVD [10].

The American Heart Association states that approximately 82.6 million individuals in the U.S. presently suffer from one or more types of CVDs, establishing it as a primary factor behind mortality in both males and females [11]. Common types of CVDs include stroke, congestive heart failure, coronary heart disease, and hypertension [12, 13]. Considering the intricate nature, risk factors, inherent genetic composition, and trajectory of CVD, personalized treatment is considered indispensable [14]. Moreover, progress in genomics has significantly contributed to comprehending the molecular pathways linked to the prevalence of CVDs [3]. These advancements were propelled by next-generation sequencing (NGS), which enabled the discovery of novel genetic correlations and the capacity to assess genetic diversity among patients [15]. Recent developments in the field of genomics and bioinformatics have greatly aided in better understanding the complex nature of CVD etiology. However, the development of an AI/ML predictive engine that utilizes genetic biomarkers to assess the risk of CVD in patients is still in its early stages [16, 17, 18]. Recent studies have explored the potential of employing AI/ML algorithms on whole genome and whole exome sequencing (WES/WGS) data for statistical and prognostic analyses for a wide variety of diseases including but not limited to Crohn’s disease [19], inflammatory bowel disease [20], breast cancer [21], colon cancer [22] and Alzheimer’s disease [23].

Previously, we have created AI/ML models to investigate and identify genes associated with heart failure (HF), atrial fibrillation (AF), and other CVDs and successfully predict these diseases with high accuracy [24]. However, one of the major limitations of our and most of the other published disease specific research using AI/ML and bioinformatics approaches is the focus on genes known to be associated with disease [2, 24, 25]. In this study, we propose a new AI/ML model that adapts an innovative nexus of algorithms to predict CVDs using critical transcriptomic biomarkers determined using our comprehensive statistical analysis (**Figure 1**). Our model is trained on an AI/ML ready dataset of whole transcriptome-based gene expression and clinical data of consented individuals. We observed novel as well as known biomarkers that were associated with CVDs, relative to our previous model [24]. We demonstrate that our current model can produce accurate predictions regarding CVD diagnosis. By identifying specific biomarkers, we have unveiled a crucial set of potential indicators for the early detection of CVDs. These biomarkers provide essential clues in identifying at-risk patients before symptoms manifest, allowing for timely intervention and improved patient outcomes. With the successful implementation of our newly developed predictive engine, healthcare professionals now have access to a valuable framework that utilizes biomarker profiles to accurately identify patients at risk of CVDs.

**Figure 1.**
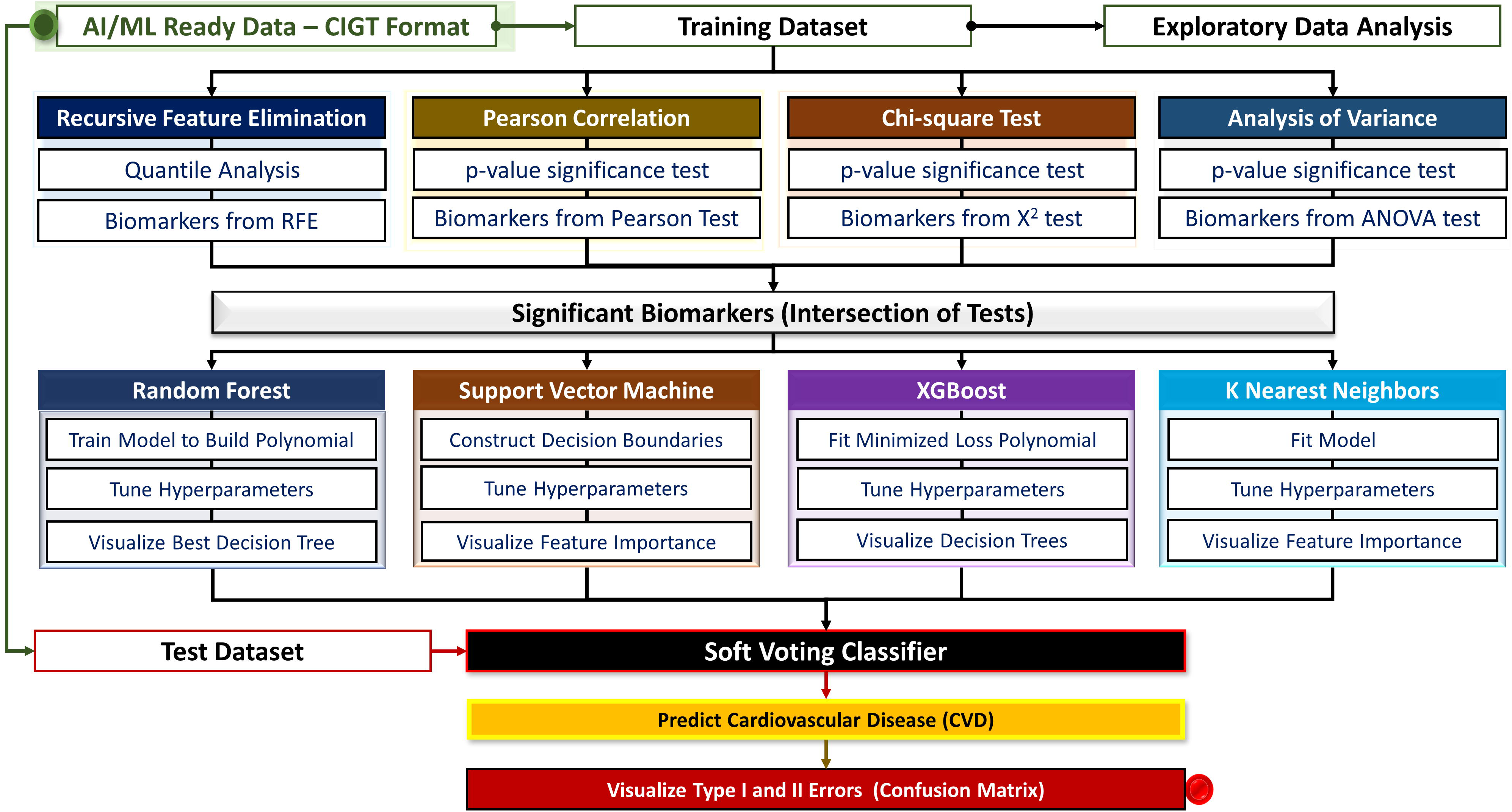
Methodology and study design, workflow, and bioinformatics. Artificial intelligence (AI) and machine learning (ML) for statistical and predictive analysis. Training dataset is used to identify significant biomarkers that are then utilized by a nexus of AI/ML algorithms to predict cardiovascular disease (CVD).

## Results

### Building Suitable Cohorts

Substantiating our approach towards discovering disease-relevant biomarkers effectively to predict patients’ diagnostic status necessitated creating a comprehensive dataset to represent our patient cohort. The cohort consisted of 61 CVD patients, including 40 males and 21 females, aged 45 to 92. The participants self-identified their race as follows: 42 were white, 7 were black or African American, 1 was Asian, and 11 were of unknown race. These individuals were clinically diagnosed with CVDs, specifically Heart Failure (HF), and Atrial Fibrillation (AF). In addition, we constructed a control group comprising 10 healthy individuals, evenly split between males and females. Among them, 9 identified as white, and 1 did not disclose their race. The age range of this group was 28 to 78 years. All procedures involving human participants were in accordance with the ethical standards of the institution and with the 1964 Helsinki Declaration and its later amendments or comparable ethical standards. All human samples were used in accordance with relevant guidelines and regulations, and all experimental protocols were approved by the Institutional Review Board (IRB) of Rutgers. Utilizing our proposed Clinically Integrated Genomic and Transcriptomic (CIGT) format, we integrated transcriptomics, clinical, and demographics data of each patient. Data pre-processing increased our cohort’s strength through the elimination of non-ubiquitous patient attributes; features present in 80% of the cohort were included and the less occurring were eliminated from the CIGT dataset to avoid extrapolation from ML classifiers downstream. Resulting from this filtration, 751 transcriptomic and clinical biomarkers remained suitable. The CIGT dataset was subset into training and testing sets, with a testing size of 30%.

### Discovering Supported Biomarkers

Statistical algorithms were applied on the training dataset to retrieve highly significant biomarkers. To assess the differences in expression levels and clinical characteristics across CVD patients and healthy individuals, we employed a convergence of four statistical algorithms: I) Recursive Feature Elimination (RFE), II) Pearson Correlation, III) Chi-Square, and IV) Analysis of Variance (ANOVA) (**Figure 2**). To ascertain the statistical significance of each algorithm, we conducted a p-value significance test and recorded the obtained p-values in a list together with the raw scores generated by each algorithm. We exercised the scientific standard of 0.05 as a threshold for our statistical significance test and utilized the logarithmic function, with a base of 10, for easier interpretation.

**Figure 2.**
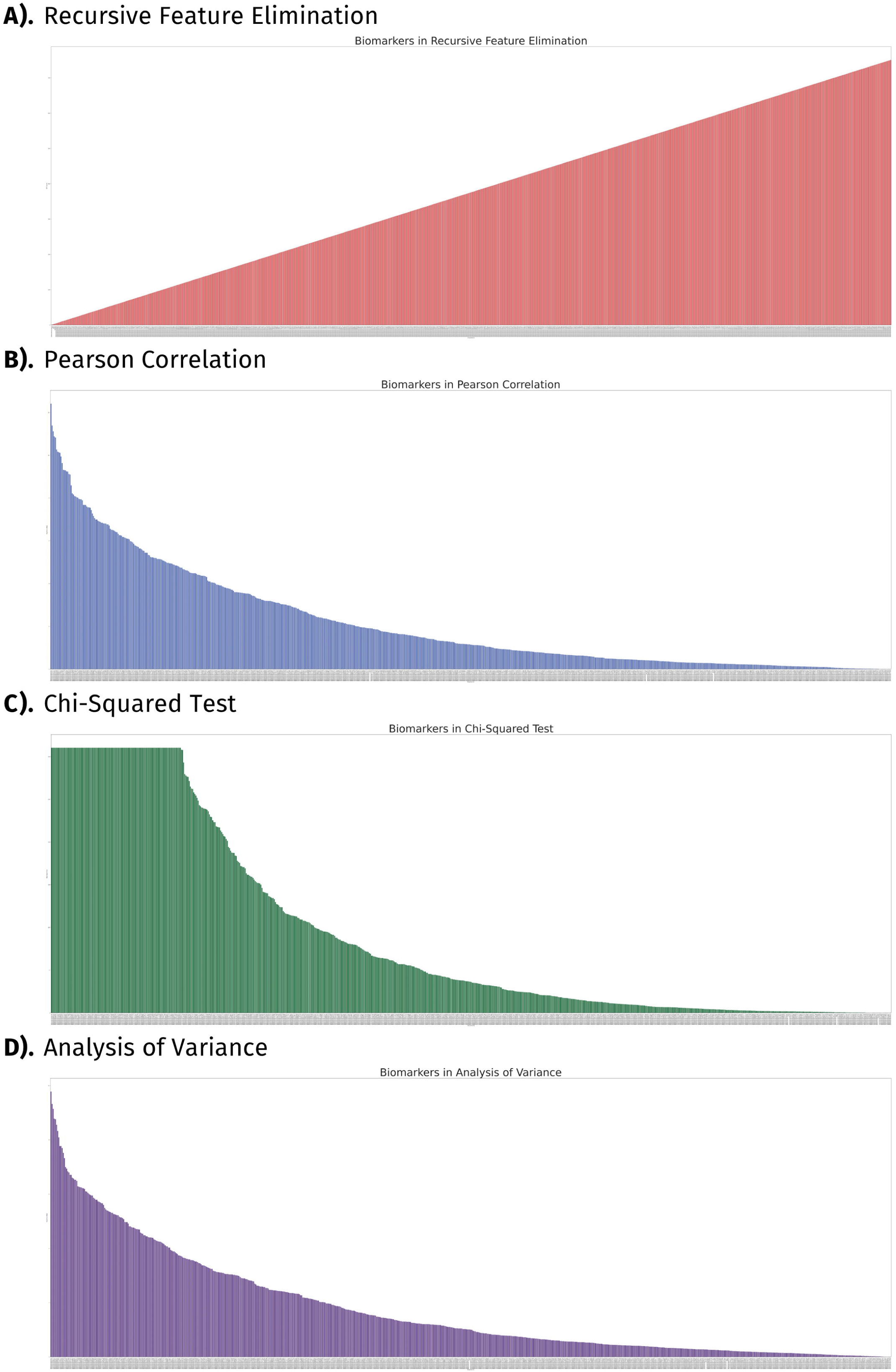
Feature Selection of Biomarkers. Statistical significance test to determine the importance of each gene according to the algorithm used. The y-axis represents the p-values as a logarithmic expression while the x-axis displays distinct biomarkers. Features are displayed from A) Recursive Feature Elimination; B) Pearson Correlation; C) Chi-Squared test; and D) Analysis of Variance.

RFE systematically eliminated the least informative features, which enabled the identification of the strongest correlations between biomarkers and CVD. The RFE algorithm assigned scores to each feature, reflecting their relative importance, with higher scores indicating lesser significance. These scores were then utilized to rank the features based on their relevance to CVD diagnosis (**Figure 2A**). Next, the Pearson correlation test was applied to quantitively assess the magnitude of linear association between biomarkers and CVD. In our study, we observed the correlation coefficient, which ranges from –1 to 1, with larger absolute values indicating a more pronounced association. However, to assess the statistical significance of the findings, we also examined the negative logarithm of the p-value for both transcriptomic and clinical features (**Figure 2B**). Notably, higher bars in the graph indicate greater significance to CVD diagnosis.

We applied the chi-square test to investigate the independence among categorical factors on CVD detection and discern any significant relationships that may exist. We calculated the chi-square statistic to quantify this independence. We utilized the ANOVA test to discern the difference in the distribution of gene expression patterns between healthy individuals and those afflicted with CVD. We computed the F-statistic to measure this variability. We found 313 biomarkers to be supported across three of our algorithms (Pearson correlation, chi-square test, and ANOVA). The presence of high outliers, such as genes *HBA1* and *HBA2*, which are beneficial in traditional selection methods but detrimental to predictive model training, diminishes importance within our RFE classifications. To counterbalance precursory approaches to subset our biomarkers, we implemented RFE. Biomarkers classified within the top 10% were endorsed for further predictive analysis (Table 1).

**Table 1.** Statistical analysis of most significant biomarkers. Table 1 includes rankings based on Recursive Feature Elimination scores, Pearson correlation, chi-square, and Analysis of Variance test. All raw scores for are included (correlation co-efficient, chi-square statistic, and f-statistic) as well as p-values that were utilized in the visualization and artificial intelligence/machine learning (AI/ML) analysis of the data.

| Ensembl ID | Recursive Feature Elimination Score | Correlation Coefficient | Pearson Correlation (p-value) | Chi-Square Statistic | Chi-Square Test (p-value) | F-Statistic | Analysis of Variance (p-value) |
| --- | --- | --- | --- | --- | --- | --- | --- |
| ENSG00000266422 | 8 | 0.573204861 | 1.42E-07 | 6099.039146 | 0 | 18.4616809 | 8.41E-05 |
| ENSG00000242574 | 27 | 0.468662916 | 3.30E-05 | 1182.198479 | 4.51E-259 | 17.33140061 | 0.000129594 |
| ENSG00000256618 | 1 | -0.498577843 | 8.30E-06 | 425.0570428 | 1.94E-94 | 15.44622846 | 0.000271697 |
| ENSG00000265150 | 10 | 0.501748748 | 7.12E-06 | 5570.193207 | 0 | 14.6231818 | 0.000378483 |
| ENSG00000234745 | 41 | 0.44430813 | 9.24E-05 | 21800.54816 | 0 | 13.25033749 | 0.000665893 |
| ENSG00000241553 | 29 | 0.437526155 | 0.000121446 | 967.241151 | 2.37E-212 | 12.82521109 | 0.000795751 |
| ENSG00000256514 | 13 | -0.422350763 | 0.000219405 | 97.15855608 | 6.40E-23 | 12.5163011 | 0.000906631 |
| ENSG00000231389 | 46 | 0.415749505 | 0.000281356 | 2762.649364 | 0 | 12.50820118 | 0.000909747 |
| ENSG00000239998 | 35 | 0.437466127 | 0.000121737 | 467.7152611 | 1.01E-103 | 11.28761072 | 0.001536451 |
| ENSG00000234741 | 42 | 0.38109307 | 0.000957704 | 250.754169 | 1.78E-56 | 10.14411903 | 0.002543671 |
| ENSG00000247596 | 20 | 0.378112312 | 0.00105766 | 169.360342 | 1.02E-38 | 10.13146467 | 0.002558096 |
| ENSG00000215845 | 66 | 0.318411748 | 0.006413323 | 324.0418477 | 1.91E-72 | 9.419225469 | 0.003526625 |
| ENSG00000269858 | 5 | 0.393315171 | 0.000631198 | 199.9036854 | 2.19E-45 | 9.331682275 | 0.003670018 |
| ENSG00000233276 | 43 | -0.38130551 | 0.000950918 | 286.051535 | 3.61E-64 | 6.823535203 | 0.011973983 |
| ENSG00000245910 | 21 | 0.290124517 | 0.013431239 | 146.3023238 | 1.11E-33 | 6.440924863 | 0.01445292 |
| ENSG00000227097 | 53 | 0.256310109 | 0.029761901 | 3696.999979 | 0 | 5.590552265 | 0.022150113 |
| ENSG00000254999 | 14 | 0.271571684 | 0.021022304 | 105.5014956 | 9.48E-25 | 5.208092813 | 0.026955423 |
| ENSG00000260592 | 11 | 0.314078232 | 0.007215015 | 45.01668698 | 1.95E-11 | 4.491244041 | 0.039268284 |

### Predicting Cardiovascular Disease

Transcriptomic attributes serve as our predictive engine’s training dataset. This engine consists of five unique classifiers to forecast case/control predictions for our testing dataset: Random Forest (RF), Support Vector Machine (SVM), Xtreme Gradient Boost (XGBoost), k-Nearest Neighbor (k-NN), and Soft Voting Classifier (SVC). Metrics, including weighted-average F1 scores and receiver operating characteristic curves (ROC), were calculated for each classifier. Weighted-average F1 scores evaluate models in circumstances where categorical predictors are not balanced. ROC-AUC provides an additional approach to ML performance evaluation, showing a probability of a binary classifier to make true predictions rather than false positives. Values approaching 1.0 in each measure suggest high performance.

RF has demonstrated practical usage within transcriptomics [25]. Optimizing RF with GridSearchCV (**Figure 4A**), using dataset-standard parameters, the decision tree classifier made the most accurate predictions. RF selected case/control correctly in 95% of testing patients. Important features involved in RF prediction include *RN7SL593P, LILRA2*, and *HLA-B* (**Figure 4A**). ROC-AUC for our RF classifier was 0.95. The weighted-average F1 score was 0.96. SVM, a classifier suited for single-diagnosis case/control predictions, performed satisfactorily. Optimized using GridSearchCV using dataset-standard parameters (Figure 4B), the SVM classifier succeeded with 91% of predictions. *MTRNR2L1, GPX1*, and *AP003419.11* are the SVM classifier’s most essential features. This model’s ROC-AUC was the highest, 0.99. The SVM classifier’s weighted-average F1 score was 0.91. XGBoost, another decision tree-based approach, provides an accessible approach to classification. The performance of XGBoost rivals our SVM classifier, scoring 91% on predictions. XGBoost was optimized with GridSearchCV using dataset-standard parameters (Figure 4C). XGBoost’s best tree functioned using *MTRNR2L1* as its sole feature. XGBoost’s ROC-AUC was 0.94. The XGBoost classifier’s weighted-average F1 score is 0.91. k-NN’s performance was feeble compared to RF, SVM, and XGBoost. Tuned with GridSearchCV using dataset-standard parameters (Figure 4D), the k-NN classifier hit 91% of predictions. This pairs with 0.85 ROC-AUC and 0.91 weighted-average F1 score. k-NN is a resource-intensive algorithm, producing worse performance at extended runtimes compared to our previous classifiers. k-NN used *MTRNR2L1, BRK1*, and *ARPC4* most when forming predictions.

RF and XGBoost classifiers proved most applicable to transcriptomic datasets. SVM performance is sufficient for case/control classifications, but diverse problems engaging multiple diseases and disorders will lead to substantial performance declines [5]. k-NN is the least appropriate for such datasets. *MTRNR2L1* was the best transcriptomic marker for CVD predictions, with top-three importance for our SVM, XGBoost, and k-NN classifiers.

### Examining Transcriptomic Predictors

Validating the detected biomarkers’ relevance to our cohort’s diagnoses necessitated an in-depth inspection of their function in prediction and prominence in previous literature. Alongside a thorough review of previous scientific findings, biomarkers correlations are reported and tied to their roles in disease classification. The literature review revealed 14 transcriptomic biomarkers linked with CVDs and a variety of other diseases and disorders within our cohort. *HLA-DMB* and *HLA-B* are associated with cardiomyopathy. *RN7SL2* and *GPX1* are associated with stroke. *ARPC4* and *LILRA2* are associated with atherosclerosis. Transcriptomic markers found within the supported list are also associated with various types of chronic diseases) and disorders (cancers, rheumatoid arthritis, and diabetes. Visualizations displaying clustered profiles of transcriptomic expression (**Figure 3B**) and their associations with biomarker’s intercorrelation (**Figure 3C**) indicate the mechanisms of disease classification. This correlation metric was supported using literature as well. Genes *TWF2* and *ARPC4* scored perfect correlations.

**Figure 3.**
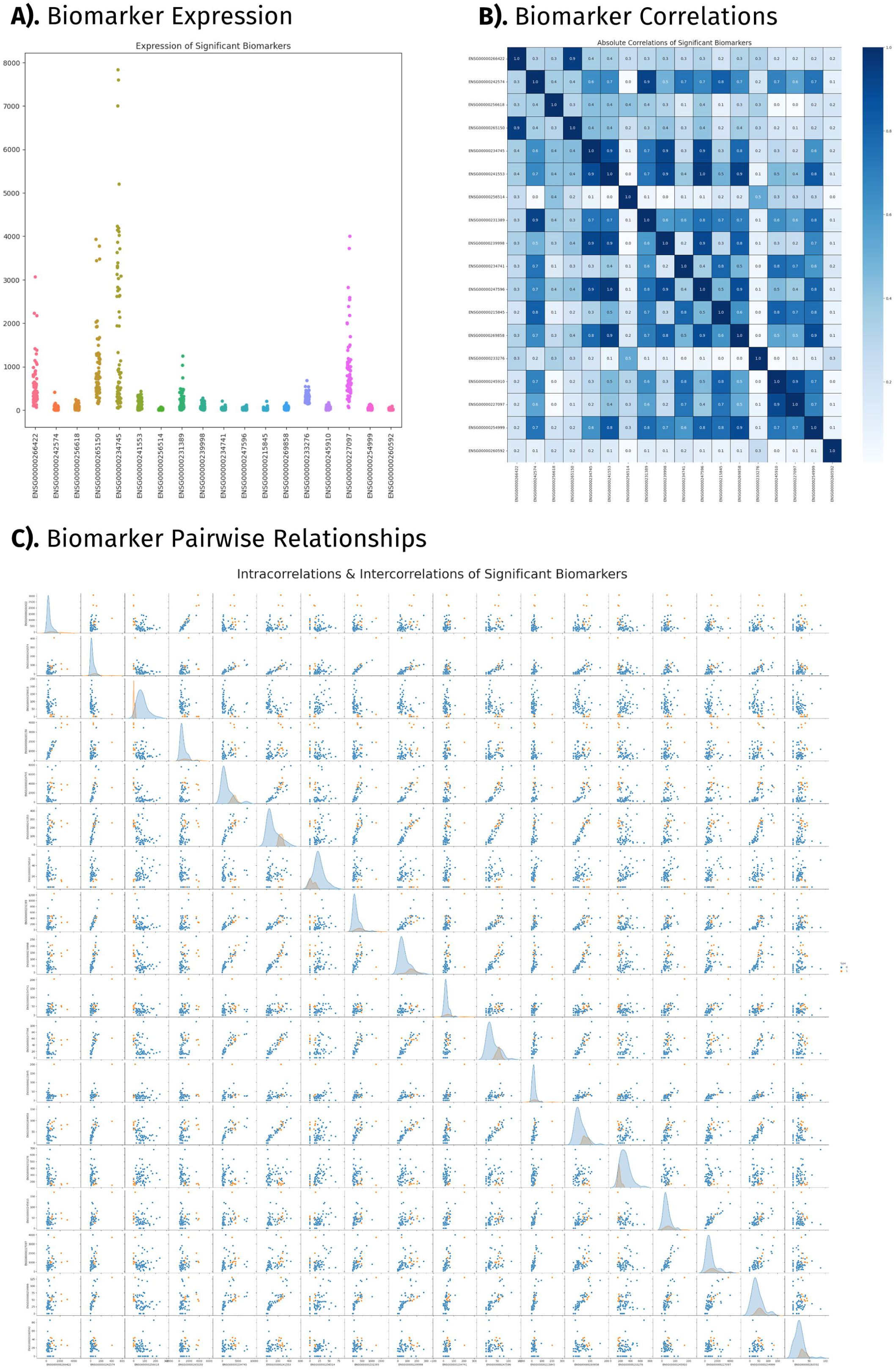
Significant biomarkers supported by four statistical tests. Results of the artificial intelligence (AI) and machine learning (ML) statistical analysis using clinically integrated genomics and transcriptomic data (CIGT) of patients with CVDs. Results include A) Biomarker expression; B) Biomarker correlations; and C) Biomarker pairwise relationships.

**Figure 4.**
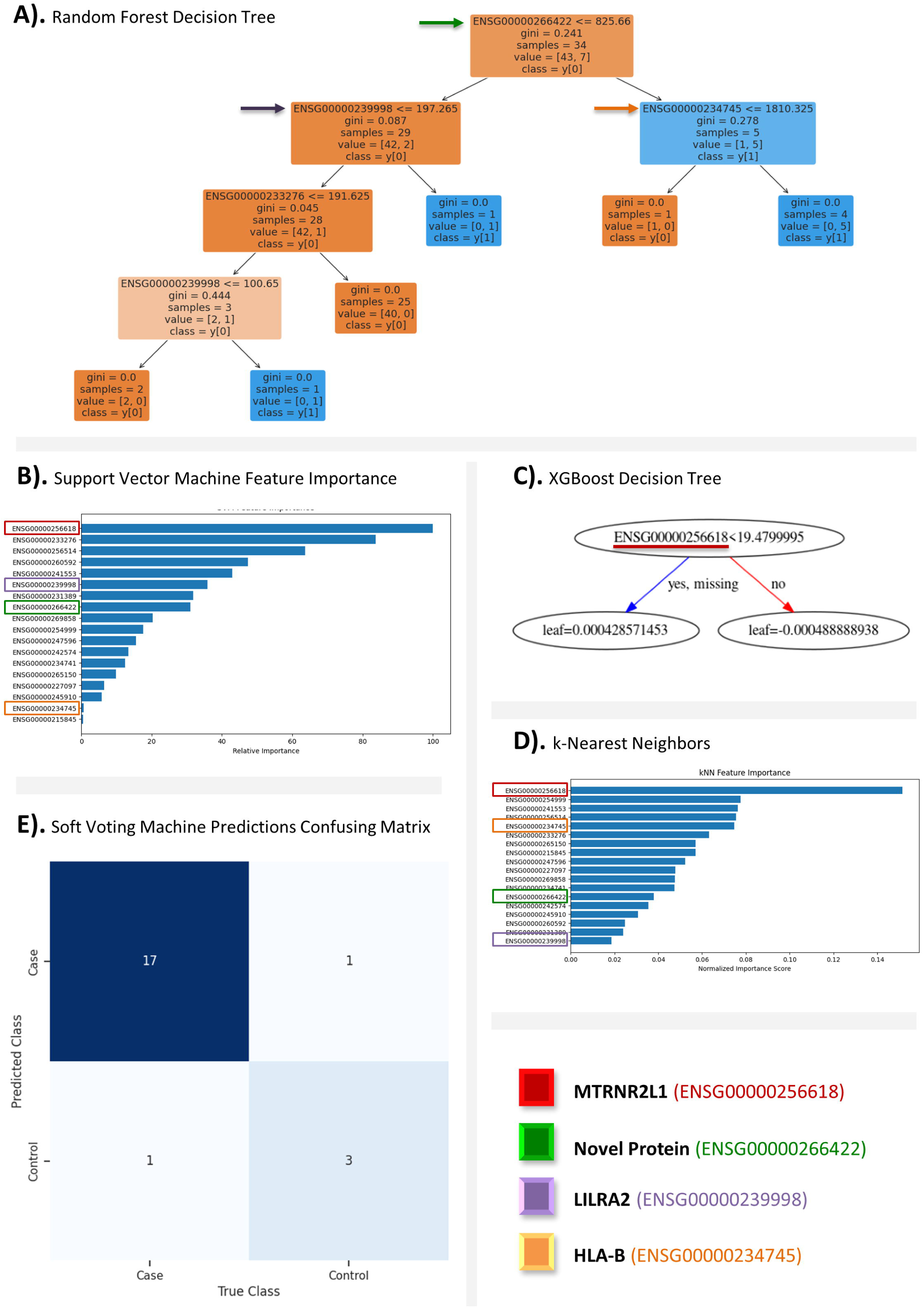
Predictive ability of our model supported by a nexus of four algorithms. Results of artificial intelligence (AI) and machine learning (ML) based predictive analysis using supported biomarkers of high-risk patients with cardiovascular diseases (CVDs). Results include A) Random Forest decision tree; B) Support vector machine feature importance; C) XGBoost decision tree; D) k-Nearest neighbors; and E) soft voting classifier predictions confusing matrix. 4 distinct biomarkers that were found to be significant within all algorithms are highlighted in the figure.

Pseudogene *MTRNR2L1* was the observed feature in all three classifiers: SVM, XGBoost, and k-NN. *MTRNR2L1* presented fluctuating expression across patients and failed to surpass a correlation above 0.5 with other transcriptomic biomarkers. *GPX1, AP003419.11*, and *CTA-363E6.6* were the three most important features of the SVM classifier beside the previously mentioned *MTRNR2L1*. *MTRNR2L1* and *GPX1* have been linked to CVDs, while *AP003419.11* and *CTA-363E6.6* have not been previously reported. These three transcriptomic markers are the least correlated with each other, the most independent function biomarkers within our list. The SVM classifier, more than others, is reliant upon independent-acting transcriptomic factors whose expression is not tied to that of another biomarker within the selected list. A cluster of highly correlated biomarkers identified, *RPS28P7, SNHG6*, and *TSTD1*, did not perform well with SVM classifier. The k-NN classifier did not display similar patterns regarding the correlation of transcriptomic biomarkers.

The XGboost classifier was reliant solely on *MTRNR2L1*, indicating the strongest association to CVDs of any transcriptomic biomarker. This algorithm ties the under expression of the lncRNA with CVD status. The RF classifier relied most prominently on the *RN7SL593P* biomarker, classifying patients below the threshold of 825.66 TPM as CVD cases. The overexpression of *RN7SL593P* has been linked to normal platelet function, a non-direct implication with CVDs. The RF classifier also placed heavy importance on *LILRA2, HLA-B,* and *GPX1* with direct links to CVDs. The decision tree algorithms contained only elements previously associated with CVDs within their optimized tree using our hyperparameter tuning metrics.

*MTRNR2L1, RN7SL593P, LILRA2*, and *HLA-B* showed the most distinct variety in their importance throughout the different classifiers. *MTRNR2L1*, scored the most important across three classifiers, but was not found in RF’s decision tree. *LILRA2* and *HLA-B* scored a correlation of 0.9, near perfect. *HLA-B* ranked as the fifth most important feature in our k-NN classifier and the second least important in the SVM classifier. *LILRA2* placed as the sixth most important feature in our SVM classifier and last in our k-NN classifier. *RN7SL593P,* the workhorse of random forest, served average throughout the remaining classifiers. These incongruencies are algorithmically dependent but may offer some understanding of underlying biological interactions between these biomarkers and CVD.

## Discussion

A persistent challenge in genomic data analysis lies in the handling and integration of large volumes of sequencing data [26]. With the implementation of our novel CIGT AI/ML ready dataset, we have begun to make significant progress to standardize heterogenous data types (genomic and clinical) for more accurate and reliable data analysis and interpretation [27]. Our novel AI/ML methodology uncovered eighteen transcriptomic biomarkers to be linked to CVDs, three of which were novel (*RN7SL593P, AP003419.11*, and *CTA-363E6.6*) and will require further analysis to understand the correlation between them and disease etiology. To further investigate gene-disease relationships for these significant biomarkers, we performed a literature review correlating these genes to CVDs (**Figure 5**).

**Figure 5.**
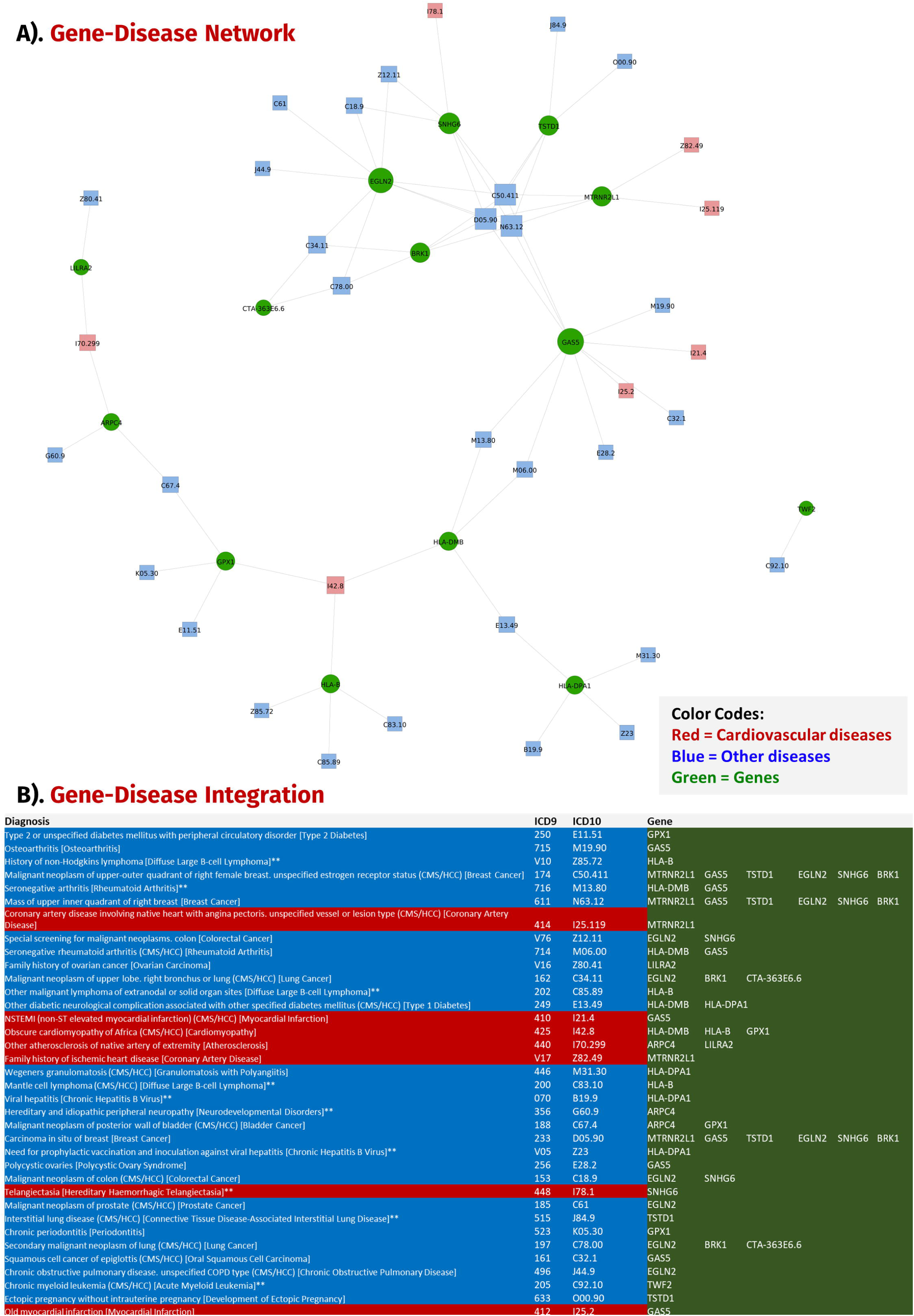
Gene-Disease network based on comparative literature review and patient diagnosis. Data validation from our cohort’s clinical records and literature review linking the supported transcriptomic biomarkers to different cardiovascular diseases (CVD) and non-CVDs. ICD-9 and 10 codes were also linked to the diagnosis list and was utilized in the network. Green corresponds to gene names; blue represents non-CVDs while red signifies CVDs.

Genes such as *HLA-DMB* [28], *HLA-B* [29], and *GPX1* [30] were found to be profoundly expressed in cardiomyopathy. While other biomarkers such as *RN7SL2* [31], *LILRA2* [32], *GAS5* [33], *TWF2* [34], *EGLN2* [35], *SNHG6* [36, 37, 38], and *BRK1* [39] have all been previously associated with phenotypic variations linked to CVD, there is limited literature associating protein-coding genes such as *RPS28P7 and CTA-363E6.6* to other known CVDs. No direct links were recorded between *RN7SL593P* and *AP003419.11* and known CVDs as well as other non-CVD-related diseases. Additional validation of these biomarkers was conducted utilizing the patients’ clinical records to elaborate on the associations between secondary diseases and their possible effect on CVD prognosis.

A significant number of biomarkers were associated to other diseases diagnosed reported for CVD patients’ clinical records. We created a network of overlapping diseases linked to the eighteen biomarkers in the highly diagnosed conditions from EHRs (Electronic Health Records) as well as those reported earlier in our comparative review (**Figure 5**). We observed that most genes were interconnected through a CVD including but not limited to cardiomyopathy, stroke, and atherosclerosis. The most common non-CVD diagnosis within our patient cohort was breast cancer, and we found *GAS5* [40], *TSTD1* [41], *EGLN2* [42], *SNHG6* [43], *BRK1* [44], and *MTRNR2L1* [45] to be indicative biomarkers. As stated earlier, cardiomyopathy was the next prevalent disease in our network corroborating our claims that our innovative AI/ML model can accurately predict CVDs. Other diseases that were shared between the genes included coronary artery disease, myocardial infarctions, lung cancer, and type 1 diabetes among others (**Figure 5**).

We believe that synergistic use of multiple AI algorithms provides more accurate results, draws insightful conclusions, and precise predictions about real-world problems compared to single AI algorithm on its own. This approach combines the best aspects of multiple machine learning algorithms into a single model. A limitation of our current study is that experimental validation is needed to support the outcomes of our AI/ML model. We addressed this constraint by utilizing clinical records and comparative literature to support our findings. In the future, our AI/ML model can be implemented in the clinical setting to aid in early disease diagnosis and improve prognosis. Our model has the potential to be generalized to investigate non-CVDs with intricate characteristics such as breast cancer, diabetes, and Alzheimer’s disease among many others. To foster these downstream applications, we have made source code openly available and freely accessible. This cutting-edge technology enhances the precision of diagnoses and empowers clinicians to tailor personalized treatment plans, ultimately leading to more effective and targeted healthcare interventions. Our findings validate the effectiveness and reliability of the model in the medical domain, offering promising prospects for improved healthcare outcomes.

### Material and Methods

Our study is divided into two major steps: I) exploratory data analysis and identification of significant biomarkers, and II) implementation of nexus AI/ML models for predictive analysis.

#### Exploratory Data Analysis and Identification of Significant Biomarkers

Exploratory data analysis was performed to show the difference in gender distribution based on the age of the subject. The box plot was presented to show a quick statistical summary (i.e., mean, median, range, outliers) of each gene in the dataset [46] as well as to explore distributions, outliers, and anomalies [46]. We utilized a convergence of statistical algorithms to evaluate the variations in expression levels and clinical characteristics between individuals with CVDs and those that are healthy. The proposed feature selection model uses four distinct algorithms: I) Recursive Feature Elimination (RFE) [47], II) Pearson Correlation [48], III) Chi-Square Test [49], and IV) Analysis of Variance (ANOVA) [50]. A combination of these tests allows the model to adapt to different matrix sizes, distributions, and attributes. All these algorithms used our CIGT dataset to compute the statistical significance of supported biomarkers by means of a p-value significance test.

To eliminate biomarkers that do not have high significance to CVD and reduce the computational load for the analysis downstream, we applied the RFE algorithm [51]. In our study, we chose the scoring metric to be based on decision trees with top 10% number of features to be from the original list of biomarkers. The correlation coefficient plays a crucial role in ranking: the higher the coefficient, the higher the rank assigned to the gene, implying a stronger association between the gene and CVD. It is important to note that a higher rank corresponds to a lower integer value. To determine each biomarker’s linear relationship to disease, we applied the Pearson correlation test where each biomarker was assigned a correlation coefficient. Subsequently, to examine the dependence between the test variable and the significant biomarkers, we employed the chi-square test. The chi-squared test has been applied widely in genomics for feature selection due to its application in multi-disease classification for genome-wide association studies (GWAS) [52]. This function requires two parameters: a scoring metric (in this case, chi-square) and ‘k’, which we set as ten. Next, we implement the ANOVA procedure, which uses a five-step approach to compute a f-statistic that determines the significance of a biomarker.

There are documented limitations associated with each testing algorithm utilized in our study. To address these challenges, we have merged these algorithms to satisfy different requirements. RFE cannot quantify the correlation between biomarkers and lacks the ability to compute multivariate significance. Furthermore, due to its iterative nature, RFE has a high time complexity [48]. One of the main limitations of the Pearson correlation test is the sensitivity to range differences between the biomarkers and their relation to disease. However, we have accounted for this by increasing the volume of data to reduce range differences between biomarkers. The main challenge associated with the chi-square test is the number of Type I and II errors in small sample sizes. However, the rationale for implementing this algorithm was to make our overall system predict better in larger matrix sizes. One more challenge that arises with ANOVA testing is the fact that if two groups of samples are of different sizes, then there is a direct issue with the strength and validity of the test. Due to the inclusion of all the other algorithms that can handle imbalances in sample size, this limitation is not of concern to this study. In our merged function, we select the statistically significant biomarkers for the ANOVA, chi-square, and Pearson correlation test and show up in the top 10% of significant biomarkers in RFE.

#### Implementation of a nexus AI/ML models for predictive analysis

The biomarkers selected were predictive for patient diagnosis and classification. We selected four algorithms for this task: Random Forest (RF) [53], Support Vector Machine (SVM) [54], K-nearest neighbors (k-NN) [55], and Extreme Gradient Boosting Decision Trees (XGBoost) [56]. We applied hyperparameter tuning to all algorithms, which were then ensembled using a Soft Voting Classifier to curate a powerful predictive engine that can perform accurate classification specific to user-specified matrices.

We started with RF, which is a meta-classifier that combines the output of multiple decision trees to categorize individuals based on their disease state. The algorithm computes a decision tree to classify patients based on their biomarker profile. The best decision tree from the forest was considered which highlights the decision boundary (i.e., polynomial) that the algorithm uses to sort patients. To classify patients based on their biomarker profile, we implemented SVM that computes support vectors. The most important classification feature highlights the relative significance of each biomarker. To further classify patients based on their biomarker profile and address limitations associated with SVM, we used the XGBoost algorithm. This algorithm computes a decision tree to highlight biomarkers that were of significance in the classification process. Finally, we applied the k-NN algorithm to determine the classification of a datapoint by majority voting amongst its ‘k’ nearest neighbors. The k-value was chosen based on iterating through all possible values of k and selecting the model with the highest accuracy.

Employing this nexus of ML algorithms helped us in navigating shortcomings that might arise from individual algorithms. The main limitation of SVMs is their inability to perform well when the data set is large [54]. However, through a combination of algorithms, SVMs can be an integral part of an ML system when the input set is small. Another limitation arises in the implementation of XGBoost where the performance is greatly diminished on sparse and unstructured data [56]. However, due to our robust data pre-processing function, we have been able to address this issue. The main limitation of k-NN is the sensitivity to feature scaling [55]. KNN calculates distances between instances to determine their similarity. If features have different scales, those with larger values can dominate the distance calculation, leading to biased results. It is essential to normalize or scale the features appropriately before applying KNN. However, KNN can adapt to changes in the training data without requiring complete retraining of the model, which is why it was selected for our analysis.

All four algorithms were ensembled using the Soft Voting Classifier, the class with the highest average probability of success was chosen as the final prediction. By combining each algorithm in this manner, the positives are accentuated while neutralizing the downsides for each algorithm.

## Acknowledgments

We appreciate great support by the Department of Medicine, Rutgers Robert Wood Johnson Medical School (RWJMS); Rutgers Institute for Health, Health Care Policy, and Aging Research (IFH); Rutgers Biomedical and Health Sciences (RBHS), at the Rutgers, The State University of New Jersey.

We thank members and collaborators of Ahmed Lab at Rutgers (RWJMS and IFH) for their support, participation, and contribution to this study.

## Authors’ information

WD is the Research Assistants at the Ahmed lab, Rutgers IFH/RWJMS.

HA is the Senior Research Assistants at the Ahmed lab, Rutgers IFH/RWJMS.

KP is the Research Assistants at the Ahmed lab, Rutgers IFH/RWJMS.

DM is the lead software engineer at Rutgers IFH.

SZ is the visiting post-doctoral researcher at Rutgers CINJ

ZA is the Assistant Professor at the Department of Medicine / Cardiovascular Disease and Hypertension, Division of General Internal Medicine, Rutgers Robert Wood Johnson Medical School, which is the part of Rutgers Biomedical and Health Sciences. Dr. Ahmed is a Core Faculty Member at the Rutgers Institute for Health, Health Care Policy and Aging Research, at Rutgers, The State University of New Jersey.

## Author contributions

ZA designed and led this study. ZA participated in sample collection, cohort building, and RNA-seq data generation. ZA performed processing, quality checking, and gene-disease data annotation and expression analysis. ZA generated AI/ML ready dataset and supported WD in designing methodology and implementing AI/ML techniques. WD, HA, DM, and SZ supported the pre- and post-computational analysis, evaluation of results. HA and ZA drafted the manuscript. All authors have participated in writing and review and have approved it for publication.

## Ethics declarations

### Ethical approval and consent to participate

Informed consent was obtained from all subjects. All human samples were used in accordance with relevant guidelines and regulations, and all experimental protocols were approved by the Institutional Review Board.

